# High-Risk Human Papillomavirus Oncogenes Disrupt the Fanconi Anemia DNA repair Pathway by Impairing Localization and Deubiquitination of FancD2

**DOI:** 10.1101/457176

**Authors:** Sujita Khanal, Denise A. Galloway

## Abstract

Persistent expression of high-risk HPV oncogenes is necessary for the development of anogenital and oropharyngeal cancers. Here, we show that E6/E7 expressing cells are hypersensitive to DNA crosslinking agent cisplatin and have defects in repairing DNA interstrand crosslinks (ICL). Importantly, we elucidate how E6/E7 attenuate the Fanconi anemia (FA) DNA crosslink repair pathway. Though E6/E7 activated the pathway by increasing FancD2 monoubiquitination and foci formation, they inhibited the completion of the repair by multiple mechanisms. E6/E7 impaired FancD2 colocalization with double-strand breaks (DSB), which subsequently hindered the recruitment of downstream protein Rad51 to DSB in E6 cells. Further, E6 expression caused delayed FancD2 de-ubiquitination, an important process for effective ICL repair. Delayed FancD2 de-ubiquitination was attributed to the increased chromatin retention of FancD2 hindering USP1 de-ubiquitinating activity, and persistently activated ATR/CHK-1/pS565 FancI signaling. E6 mediated p53 degradation did not hamper the cell cycle specific process of FancD2 modifications but abrogated repair by disrupting FancD2 de-ubiquitination. Further, E6 reduced the expression and foci formation of Palb2, which is a repair protein downstream of FancD2. These findings uncover unique mechanisms by which HPV oncogenes contribute to genomic instability and the response to cisplatin therapies.

**AUTHOR SUMMARY:** High-risk human papillomavirus (HPV) causes nearly all cervical and many other anogenital cancers, and oropharyngeal cancers. As cisplatin is the most commonly used drug for cervical and HPV-associated oropharyngeal cancers, it is important to understand how HPV oncogenes disrupt the Fanconi anemia (FA) pathway involved primarily in the repair of cisplatin-induced DNA crosslinks. However, the mechanism by which HPV E6 and E7 attenuate the FA pathway is poorly understood. We demonstrate that E6/E7 expression disrupts crosslink repair and increase cisplatin sensitivity, and attenuate the FA pathway through multiple unique mechanisms. First, E6/E7 causes accumulation of FancD2, a central component of the FA pathway, at the sites away from DNA damage. This results in reduced recruitment of Rad51, another repair protein involved in the pathway. Second, E6 causes delayed FancD2 de-ubiquitination, an important process for effective repair. Third, E6 expressing cells decreases the expression and foci formation of Palb2 repair protein. Together, this work elucidates the mechanisms by which HPV attenuates the repair of DNA crosslinks increasing cisplatin cytotoxicity and efficacy in treating HPV-positive cancers.

## INTRODUCTION

High-risk human papillomavirus (HR-HPV) E6/E7 oncoproteins are essential for the development of malignancies of the anogenital tract and oropharynx, with HPV16 being the predominant type (1). Cervical and oropharyngeal cancers are the most common HPV-associated malignancies among females and males, respectively (2). Persistent HPV infection destabilizes the cellular genome which can lead to cancer. Genomic instability is likely the result of the numerous interactions of HPV oncoproteins with host tumor suppressors and DNA damage repair (DDR) proteins. Recently, we demonstrated that high-risk HPV oncogenes attenuate double-strand break (DSB) repair by impairing the homologous recombination pathway (3). To further elucidate the mechanisms by which HPV oncogenes impair DDR, the present study focuses on the impact of HPV16 oncogenes on the Fanconi anemia-BRCA (FA or FA-BRCA) pathway.

The FA-BRCA pathway is involved in the repair of intra or interstrand crosslinks (ICL), thereby maintaining genomic stability (4, 5) (Fig S1). ICLs block DNA replication and transcription and, thus, are highly cytotoxic to cells if not repaired. The FA pathway is composed of 22 FA proteins (identified to date) (6). When any one of the FA genes of the FA-BRCA pathway is mutated, individuals have a spectrum of disorders, called Fanconi Anemia, characterized by bone marrow failure, congenital malformations, cancer predisposition, and cellular sensitivity to ICL-inducing agents (7). Upon exposure to DNA crosslinking agents, and during S-phase of the cell cycle, ATR/CHK1 signaling gets activated and helps in the formation of an ubiquitin ligase complex (FA core complex), which is composed of eight FA proteins (FancA, -B, -C, -E, -F, -G, -L, and -M) with other associated proteins (such as FAAP-24, MHF2). Among the FA core complex subunits, FancM recognizes the stalled replication fork at the ICL site and forms a landing platform for the core complex (8). Activated ATR/CHK1 phosphorylates several FA proteins including FancM, FancD2, and FancI (9). The FA core complex through its E3 ubiquitin ligase subunit FANCL and the corresponding E2 ubiquitin-conjugating enzyme (UBE2T/FancT) catalyzes the monoubiquitination of the FancD2-FancI heterodimer. Monoubiquitinated FancD2 (FancD2-Ub) and FancI-Ub form discrete nuclear foci at double-strand breaks (DSB) created at ICL-stalled replication forks, and subsequently recruit downstream DNA repair proteins, including FancD1 (BRCA2), FancS (BRCA1), FancR (Rad51), FancN (Palb2), and FancJ (BRIP1). These proteins in nuclear foci co-operate with other DNA repair pathways such as nucleotide excision repair, homology recombination, and translesion synthesis to repair ICLs (8). Once DNA is repaired, the FA pathway is turned off by de-ubiquitination of the FancD2/FancI complex to prevent prolonged cell cycle arrest and cell death (10). FancD2/FancI de-ubiquitination is catalyzed by the ubiquitin-specific protease USP1, in conjunction with UAF1 (USP1-associated factor 1). While monoubiquitination of FancD2 is essential for ICL repair, its deubiquitination by the USP1-UAF1 complex is also critical for a functional FA pathway (10-14). Knockout of either USP1 or UAF1 in mice causes an FA-like phenotype (11, 12, 14) and USP1 disruption or the absence of de-ubiquitination abrogates FancD2 foci formation and ICL repair and increases sensitivity to DNA cross-linkers (10-13).

Several studies have documented an interaction between HPV and the FA pathway. First, loss of FancA or FancD2 lead to hyperproliferation of HPV+ hyperplasia and increased proliferation of HPV genomes in organotypic cultures (15, 16). Loss of FancD2 potentiates E7 driven cancers of the female lower reproductive tract, and head and neck in two separate studies using mouse models (17, 18). Second, FA patients are susceptible to oral and anogenital squamous cell carcinomas (19), though the involvement of HPV in these cancers is controversial because of inconsistent detection of HPV DNA (20-23). Third, HPV+ head and neck cancer cell lines show greater sensitivity to cisplatin compared to HPV negative cells (24).

The molecular mechanism(s) by which HPV interacts with the FA pathway is not well-understood. HR-HPV (mainly E7) was shown to upregulate several FA genes and activate the FA pathway by increasing FancD2 monoubiquitination and foci formation (25-27). In contrast, HPV oncogenes were reported to perturb the functions of several FA proteins, including ATR, BRCA2/FancD1, BRCA1/FancS, and Rad51/FancR (3, 28-30). Because HPV increases sensitivity to DNA crosslinkers (24), and a functional FA pathway restricts HPV replication (16), and FancD2 is preferentially recruited to the HPV episome leaving the cellular genome unrepaired (31), we hypothesized that HPV oncogenes impair the FA DNA repair pathway to facilitate the HPV life cycle. To date, studies have shown that HPV increases monoubiquitination of FancD2/FancI (25-27, 31), but none have studied their deubiquitination pattern, which is as important as monoubiquitination for ICL repair (10-13). Our study shows that HPV E6 caused delayed deubiquitination of FancD2 though it increased FancD2 monoubiquitination. Similarly, HPV E6 and E7 expressing cells increased FancD2 foci formation but impaired FancD2 colocalization to double-strand breaks. Collectively, our data demonstrate that HPV16 oncogenes abrogate the FA pathway, which further supports the hypothesis that HPV inhibits cellular DDR causing genomic instability.

## RESULTS

### HPV16 oncogenes impair ICL repair and depend on the FA pathway for cisplatin sensitivity

We have previously shown that β-HPV 5/8 E6 and HPV16 E6 expression increases sensitivity to crosslinking agents such as cisplatin, mitomycin C and UVB (28). To determine if the expression of E7 could also increase crosslinker sensitivity, we expressed HPV16 E6 and E7 individually or together in primary human foreskin keratinocytes (HFKs). Expression of HPV16 E6 and E7 was confirmed by qRT-PCR, as well as by immunoblot for their established targets p53 and pRB (Fig S2). We found that both E6 and E7 expressing cells are hypersensitive to cisplatin compared to LXSN control (Fig 1A). When IC50 values were compared, E6 and E7 expressing cells were respectively 3.4 and 2 times more sensitive to cisplatin than LXSN.

**Fig 1.**
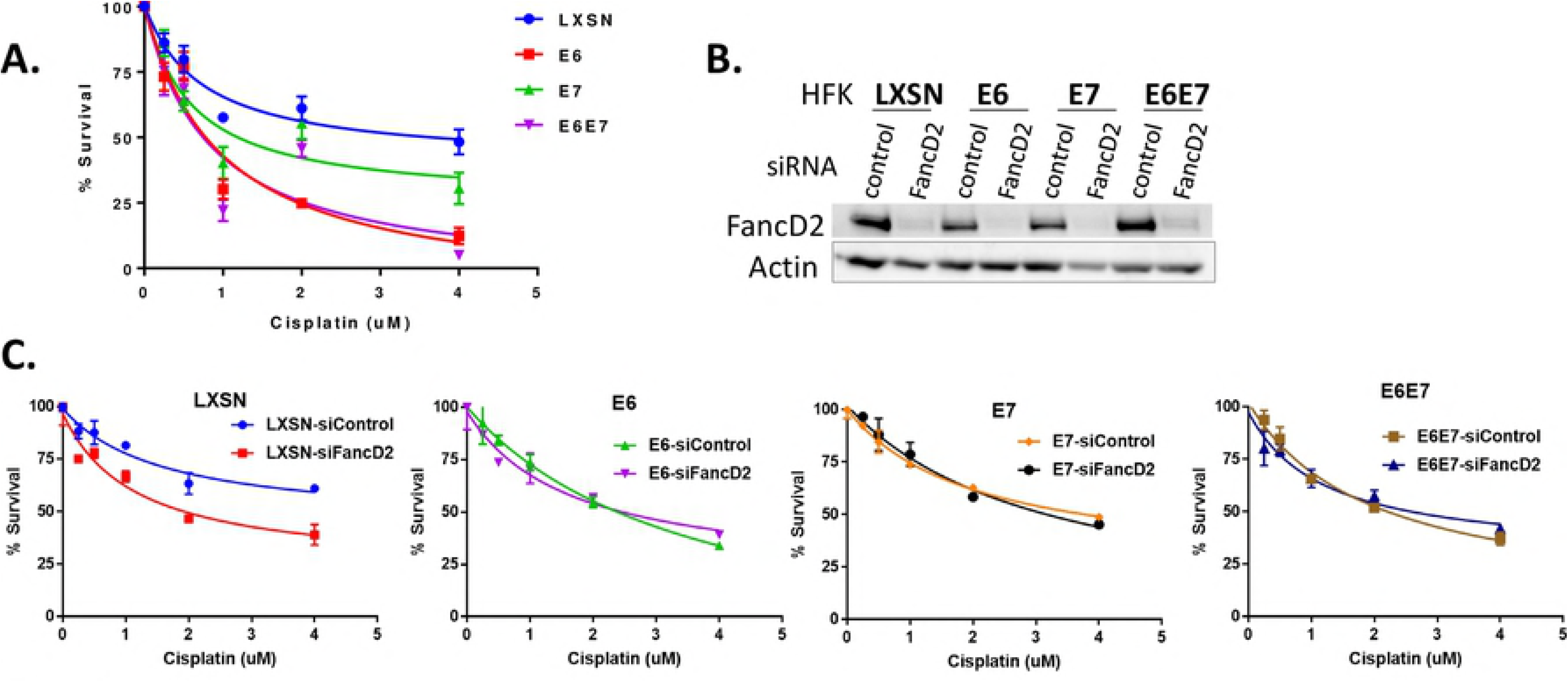
HPV oncogenes expressing cells impair ICL repair and depend on FancD2 for cisplatin sensitivity. (A) Cells were treated with increasing concentrations of cisplatin for 72 hr and % survival was measured using the Crystal Violet assay. LXSN, E6, E7, E6E7 expressing cells showed respective IC50 of 2.5, 0.74, 1.27 and 0.73 uM; 72 hr. (B-C) Cells were transfected with siControl or siFancD2 for 48hrs and plated for western blotting and cisplatin survival assay. (B) Western blot showing FancD2 knockdown in the cells harvested at the time of reading survival assay. (C) Survival curves of LXSN, E6, E7 and E6E7 cells transfected with siRNA and treated with indicated doses of cisplatin for 48 hr. (D)

Genetic epistasis analysis was conducted to determine whether the cisplatin hypersensitivity observed in E6/E7 expressing cells is due to a defect in the FA pathway. For this, we knocked down FancD2 and assessed cisplatin sensitivity (Fig 1B-C). FancD2 protein depletion by siRNA was confirmed by immunoblotting. FancD2 knockdown in LXSN control cells resulted in further increased sensitization to cisplatin, whereas there was little/no effect on cisplatin sensitivity in E6 or E7 expressing cells (Fig 1C). These results suggest that cisplatin sensitivity of HPV oncogene expressing cells is due to a defect in the FA pathway, particularly at or downstream of FancD2.

The ability of E6/E7 expressing cells to repair cisplatin-induced ICLs was investigated by utilizing a modified comet assay which has been widely used to evaluate ICL repair *in vivo* at the single cell level (32-34). Cells were treated with cisplatin for 2 hr and then incubated in cisplatin-free medium. At 24, 48 and 72 hours post-treatment, cells were harvested and frozen to analyze all samples concurrently. Cells were thawed and irradiated immediately prior to the comet assay, to deliver a fixed level of random DSBs (Fig 2A). Crosslinks hold the two strands of DNA together during alkaline denaturation and retard electrophoretic mobility of the irradiated DNA resulting in reduced tail moment compared to untreated irradiated controls. The tail moment was at a basal level in untreated and unirradiated cells (Fig 2B). At 0 hr post-cisplatin treatment, the tail moment was decreased in all cell types compared to corresponding irradiated untreated controls. At 72 hr post-cisplatin treatment, LXSN cells regained the tail moment indicating that these cells were able to repair crosslinks. In contrast, E6E7 expressing cells did not improve the tail moment 72 hr after cisplatin treatment, suggesting that ICLs remained unrepaired in these cells (Fig 2C). Further, repair kinetics of cisplatin-induced ICLs were expressed as the percentage of crosslinks remaining at the time points assessed (Fig 2D). ICLs present in the cisplatin-treated sample were calculated by comparing the tail moment of treated and irradiated (Cp.IR) cells with irradiated (IR) samples and untreated (Ø) control samples (as described in Methods). Cisplatin-induced ICLs were removed efficiently in LXSN cells with ~35% of the ICLs remaining at 72 hr, whereas in E6 or E7 expressing cells, significantly elevated levels of ICLs persisted and remained unrepaired even at 72 hr. Increased formation of % ICLs in E6/E7 cells at 24, 48 and 72 hr indicate a possible conversion of monoadducts or intrastrand adducts to interstrand crosslinks (35). Taken together, these data support the hypothesis that HPV oncogene expressing cells have a decreased ability to repair cisplatin-induced ICL.

**Fig 2.**
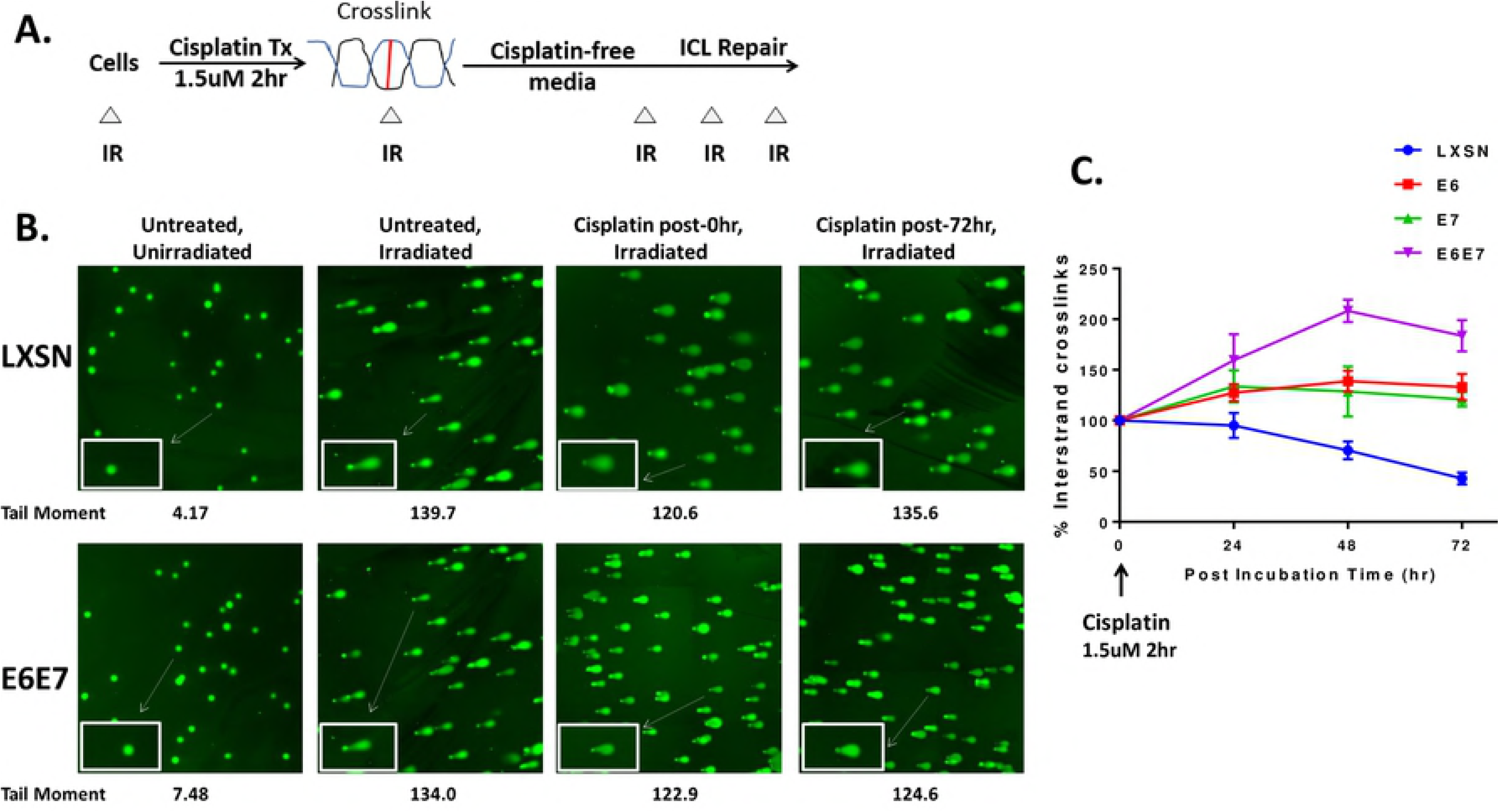
HPV oncogenes impair the repair of ICLs. (A) Outline of an experiment to evaluate repair of cisplatin-induced ICLs using the modified alkaline comet assay. Cells were treated with 1.5 μM cisplatin for 2 hr and then incubated in cisplatin-free medium. Triangles (Δ) represent the time points (0, 24, 48 and 72 hr after cisplatin release) when cells were harvested and frozen. Immediately before comet analysis, cells were thawed and irradiated with 12 Gy to deliver a fixed level of random DSBs. (B) Representative comet images of untreated and cisplatin treated+ irradiated cells at 0hr and 72 hr. The insets are the magnified comets. A mean tail moment from a representative assay is shown. (C) Quantification of % ICL present in LXSN and E6/E7 expressing HFK cells at different post-treatment incubation time. The percentage of ICLs remaining was determined as described in Methods. Data represent mean ± SEM of at least 3 independent biological replicates.

### FancD2/FancI monoubiquitination is not impaired by HPV oncogenes

To screen for defects in the FA pathway, we first examined the levels of FancD2 in HPV oncogene expressing cells. When untreated cells were analyzed, total FancD2 (Ub + non-Ub) levels were significantly increased in E6/E7 expressing cells compared to LXSN control (Fig 3A). When E6 or E7 were individually expressed, cells had about 2 times more total FancD2 compared to LXSN; but the level increased by ~4 fold when E6 and E7 were expressed together. Total FancI (Ub + non-Ub) levels were also proportionately increased in E6/E7 expressing cells compared to LXSN.

**Fig 3.**
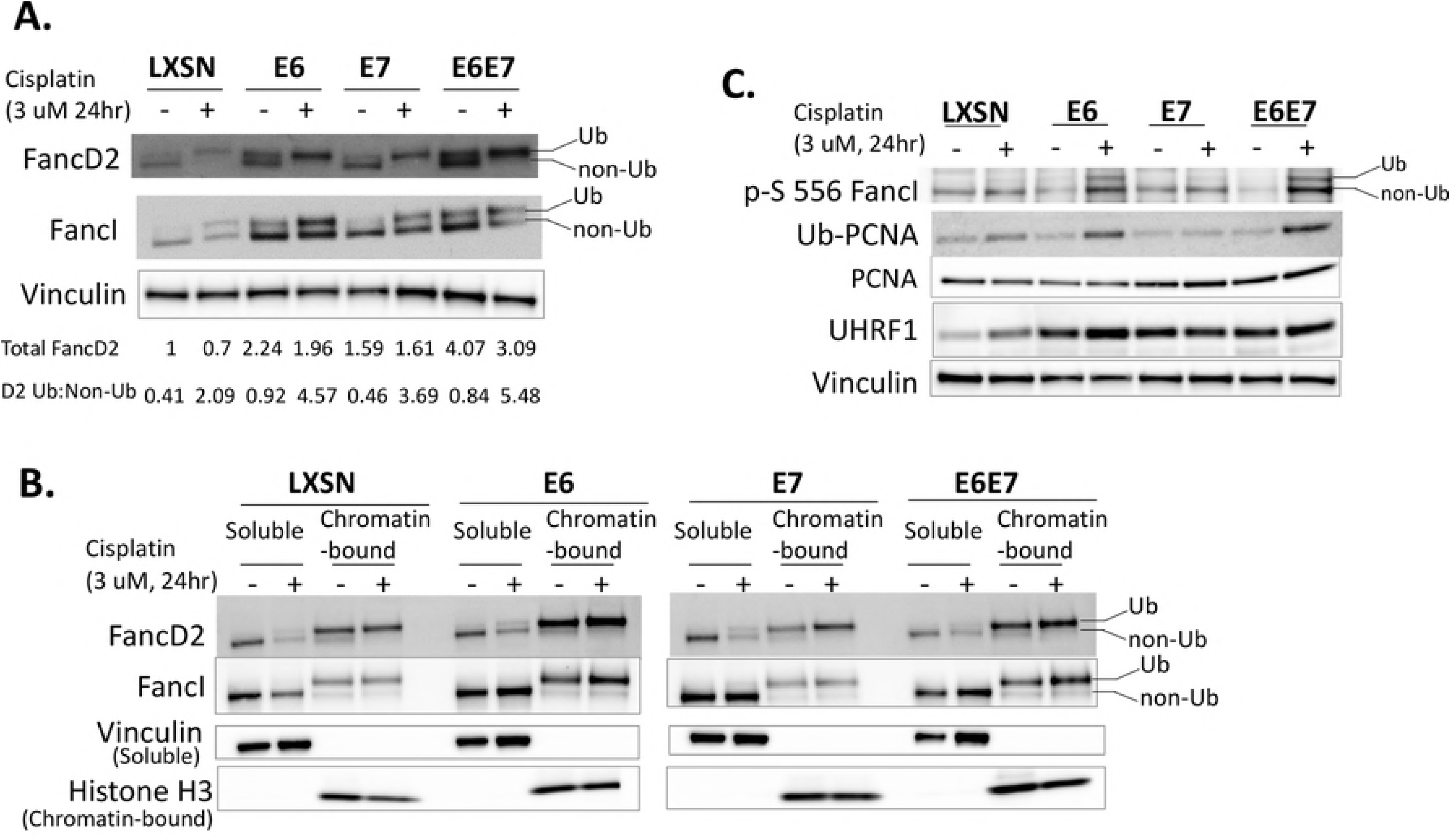
HPV oncogenes increase chromatin-bound monoubiquitinated FancD2/FancI. (A) Immunoblot showing FancD2/FancI expression and monoubiquitination status in transduced HFK cells which were either untreated or treated with 3 uM cisplatin for 24 hr. Ub refers to the monoubiquitinated forms of FancD2 and FancI, and non-Ub refers to the non-ubiquitinated forms. Ratios of monoubiquitinated to non-ubiquitinated FancD2 (D2 Ub: Non-Ub) and total FancD2 (Ub + Non-Ub) levels are indicated beneath the corresponding lanes. (B) Immunoblot of soluble and chromatin-bound fractions prepared from transduced HFK cells that were either untreated or treated with 3 uM cisplatin for 24 hr. Vinculin and Histone H3 act as loading controls respectively for soluble and chromatin-bound fractions. (C) Immunoblot of whole cell lysates showing levels of phosphorylated S556 FancI, Ub-PCNA, non-Ub PCNA, and UHFR1. Vinculin acts as a loading control.

FancD2 monoubiquitination was evaluated in E6/E7 expressing cells as a readout to define an activated FA pathway. Ub-FancD2 or Ub-FancI can be distinguished from the non-Ub form as a retarded mobility on gels. E6 and E7 increased FancD2 and FancI monoubiquitination both at baseline and after cisplatin treatment (Fig 3A). Approximately a 3-fold increase in the Ub-FancD2: non-Ub FancD2 ratio was observed in E6/E7 expressing cells on cisplatin treatment. Similar results were observed following MMC treatment and UVB irradiation (Fig S2C-D). When cells were analyzed following different lengths of cisplatin treatment, LXSN cells showed Ub-FancD2 after 6 hr of cisplatin treatment; but Ub-FancD2 was found in E6 cells even without cisplatin treatment (Fig S2E). Further, we performed cellular fractionation analyses and prepared chromatin and soluble fractions from HFKs expressing HPV oncogenes or LXSN control. As expected, Ub-FancD2 or Ub-FancI was enriched dramatically in the chromatin-bound fraction of all transduced HFK cells. Importantly, an increased recruitment of Ub-FancD2/FancI to chromatin was detected in E6 expressing cells compared to LXSN control and E7 expressing cells (Fig 3B).

We next sought to know the effectors that may contribute to increasing FancD2 monoubiquitination in E6/E7 expressing cells. Phosphorylation of FancI S556 occurs upstream of, and enhances, FancD2 monoubiquitination (7). Additionally, the proliferating cell nuclear antigen (PCNA) is monoubiquitinated by the RAD18 ubiquitin ligase in response to ICL lesions. Apart from its role in translesion synthesis repair, ubiquitinated PCNA (PCNA-Ub) is known to promote FancD2 monoubiquitination by either facilitating FancD2 recruitment onto chromatin via a direct physical interaction (36) or by promoting the recruitment of FancL and its E3 ubiquitin ligase activity on FancD2 (37, 38). Further, the protein UHRF1 (ubiquitin-like with PHD and RING finger domains 1) has recently been identified as a sensor of ICLs and is required for the recruitment of FancD2 to ICL sites (39, 40). Phosphorylated FancI-S556, Ub-PCNA, and UHFR1 levels were increased in E6 expressing cells compared to LXSN following cisplatin treatment (Fig 3C). These results suggest that elevated p-S556 FancI, Ub-PCNA and UHRF1 levels may be involved in the increased chromatin-bound fraction of Ub-FancD2 in E6 cells.

### HPV oncogenes do not disrupt FancD2 nuclear foci formation but impair colocalization of FancD2 with double-strand DNA breaks

To further investigate the interaction of HPV with the FA pathway, the ability of E6/E7 expressing cells to form nuclear foci of FancD2 was quantified as the percentage of cells with >5 FancD2 foci in HFKs. FancD2 foci formation in cisplatin-treated E6 or E7 cells was elevated compared to LXSN controls (Fig 4A-B). Even without cisplatin treatment, there was increased FancD2 foci formation in E6 expressing cells. Phospho-H2AX foci were used as markers for DNA double-strand breaks (DSB).

**Fig 4.**
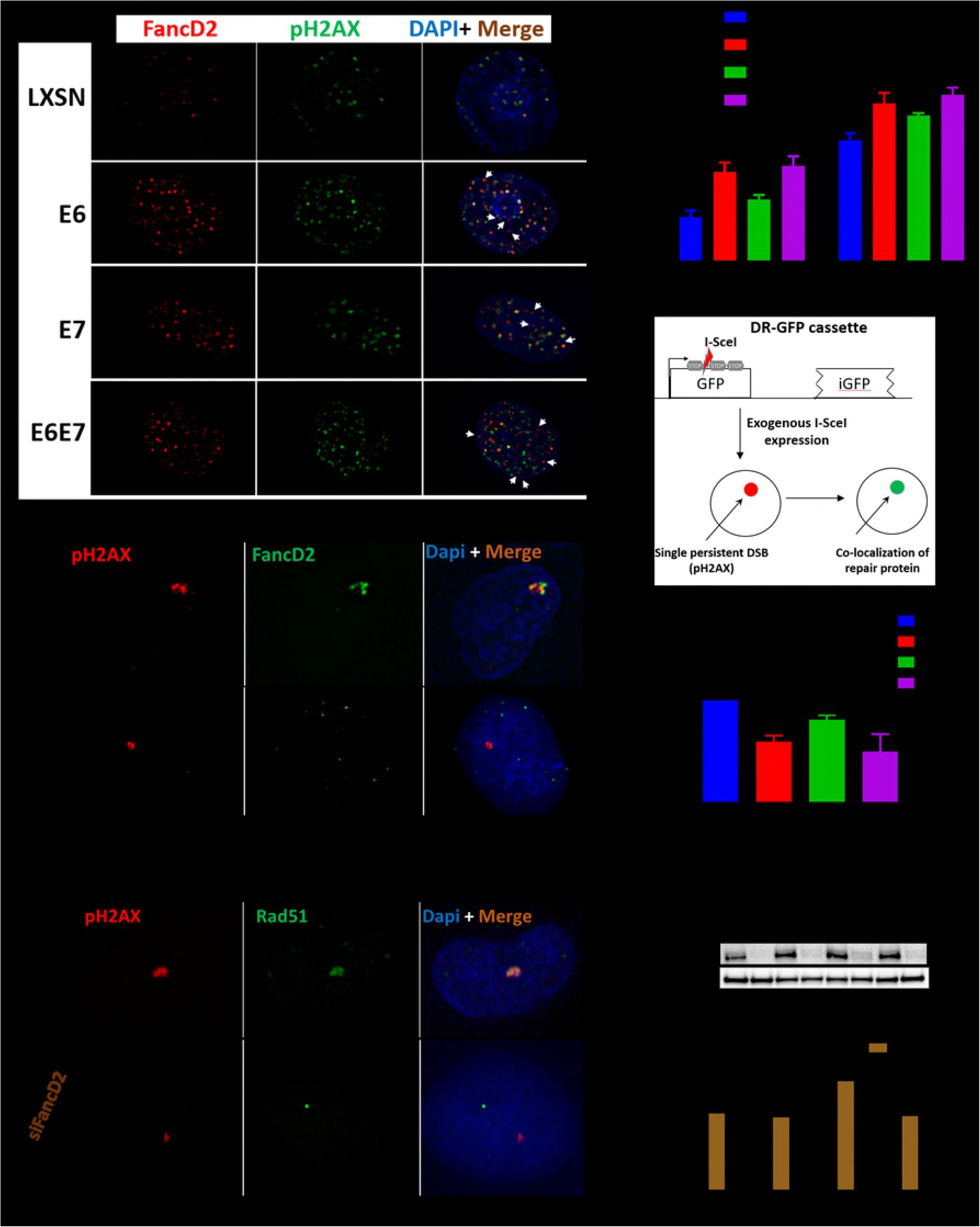
HPV oncogenes increase FancD2 foci formation but impair colocalization of FancD2 with DSBs, and mislocalized FancD2 in E6 cells causes reduction in Rad51 recruitment to DSBs. (A)HFK cells were treated with cisplatin (3 uM) for 24 hr and immunostained with FancD2 (red), pH2AX (green) and DAPI (blue). Representative images are shown. White arrows denote mislocalization of FancD2 away from pH2AX. (B) Cells with >5 foci were counted, and the percentage of positive cells is plotted (n = 3, mean ± SEM). (C) Schematic of I-SceI colocalization assay. The DR-GFP reporter cassette (integrated into U2OS genome) consists of two copies of nonfunctional GFP gene. The first copy is inactive due to the presence of a stop codon within the I-SceI cleavage site, while the second copy (iGFP) is truncated at both ends. Exogenous expression of I-SceI in U2OS cells with one integrated copy of the ISceI recognition site produces a single persistent DSB. Recruitment of repair protein (green) to this enlarged pH2AX focus (red) can be visualized by IF. (D) U2OS-DR cells (transduced with LXSN or E6/E7) were transfected with I-SceI expression plasmid for 24 hr before fixation. Cells were immunostained with pH2AX, FancD2, and DAPI (blue). Cells with a single large pH2AX focus (red) were examined for the colocalization with FancD2 (green). Representative images are shown. (E) Quantification of the frequency of colocalization of FancD2 with pH2AX foci in U2OS-DR cells. Data represent mean ± SEM and was based on observations from ≥ 50 cells from at least three independent experiments. (F-H) U2OS-DR cells transduced with the indicated constructs were transfected with the siControl or siFancD2. They were transfected with I-SceI expression plasmid and then fixed after 24 hr of transfection and stained with Rad51 and pH2AX antibodies. (F) Cell lysates were subjected to western blotting to confirm depletion of FancD2. (G) Cells with a single large pH2AX focus (red) were inspected for the colocalization with Rad51 (green). Representative images are shown. (H) Quantification of the frequency of colocalization of Rad51 with pH2AX foci. Data represent mean ± SEM and was based on observations from ≥ 25 cells from at least three independent experiments. * and ** indicate significance respectively at p<0.05 and p<0.01 (compared to LXSN) whereas n.s. indicates non-significant.

We previously reported that HPV oncogene expressing cells impair the colocalization of Rad51 with pH2AX (3). The same approach was used to investigate whether E6/E7 expressing cells affect the localization of FancD2 to DSBs. Initial experiments suggested that the expression of HPV oncogenes may cause FancD2 to be localized away from DSBs (as shown by white arrows in cisplatin-treated HFKs transduced with E6/E7, Fig 4A). However, mislocalization was difficult to quantitate in the nucleus with the numerous DSBs induced by cisplatin. For this, we utilized U2OS-DR cells transduced with LXSN, E6, E7 and E6E7 and transiently transfected with an I-SceI expression vector, as previously described (3). These U2OS cells have clonally integrated DR-GFP cassette consisting of two copies of nonfunctional GFP (Fig 4C) (41). Exogenous expression of I-SceI produces a single DSB within the first GFP gene which contains an I-SceI recognition site. Thus, cells with single large pH2AX foci were selected and inspected for its colocalization with FancD2 (Fig 4C). An excellent colocalization of FancD2 with pH2AX was observed in LXSN cells, but there was ~50% and 20% reduction in colocalization of FancD2 in E6 and E7 expressing cells, respectively compared to LXSN cells (Fig 4D-E). These data suggest that FancD2 is forming unproductive repair complexes by localizing away from DSBs in E6/E7 expressing cells.

### FancD2 is responsible for Rad51 mislocalization in E6 cells

Mislocalization of FancD2 in E6 cells (Fig 4D-E) may hinder the recruitment of downstream proteins, such as Rad51 to sites of DNA damage. In fact, several lines of evidence indicate that FancD2 helps in the localization of Rad51 to DSBs. First, Ub-FancD2 colocalizes with Rad51 in nuclear foci during S phase (42). Second, FancD2 promotes recruitment of Rad51 in nuclear foci by directly binding and stabilizing Rad51-DNA nucleoprotein filament for DSB repair (43, 44). Third, FancD2 colocalized with Rad51 in cisplatin treated HFK LXSN control cells (Fig S3). We previously reported that the colocalization of Rad51 with DSB is impaired in E6 expressing cells (3). To determine whether the mislocalization of Rad51 observed in E6 expressing cells is epistatic to FancD2, the Isce-I colocalization assay was repeated, as described above (Fig 4C), in FancD2-depleted cells and immunostained with Rad51 and pH2AX. Western blot analysis confirmed FancD2 protein depletion in the siFancD2- transfected cells compared to siControl cells (Fig 4F). FancD2 depletion in LXSN cells caused a significant reduction in colocalization of Rad51 to pH2AX in I-SceI induced DSB (Fig 4G-H), supporting the idea that FancD2 promotes Rad51 recruitment to DSB in normal cells. However, there was no significant reduction in the % Rad51 colocalization in FancD2 depleted E6 expressing cells, suggesting that the Rad51 recruitment defect associated with E6 cells is due to FancD2 and therefore cannot be reduced further with siFancD2.

### HPV E6 causes delayed de-ubiquitination of FancD2

To further screen for defects in the FA pathway, we investigated how E6/E7 affects the de-ubiquitination pattern of FancD2/FancI upon DNA repair. Though FancD2/FancI monoubiquitination is considered as a functional activator of the FA pathway, de-ubiquitination of FancD2/FancI is also critical for effective ICL repair (10-13). Since E6/E7 increases monoubiquitination of FancD2/FancI, there may be a defect in de-ubiquitination. To address this, cells were treated with cisplatin or exposed to UV and allowed to recover for the indicated time (Fig 5A). Delayed de-ubiquitination of FancD2 was observed in E6 expressing cells during recovery after UVB exposure or cisplatin removal (Fig 5A-C). Most of FancD2 was de-ubiquitinated following 24 hr of UVB and cisplatin release in LXSN cells but not in E6 or E6+E7 expressing cells. E7 expressing cells behaved more like LXSN.

**Fig 5.**
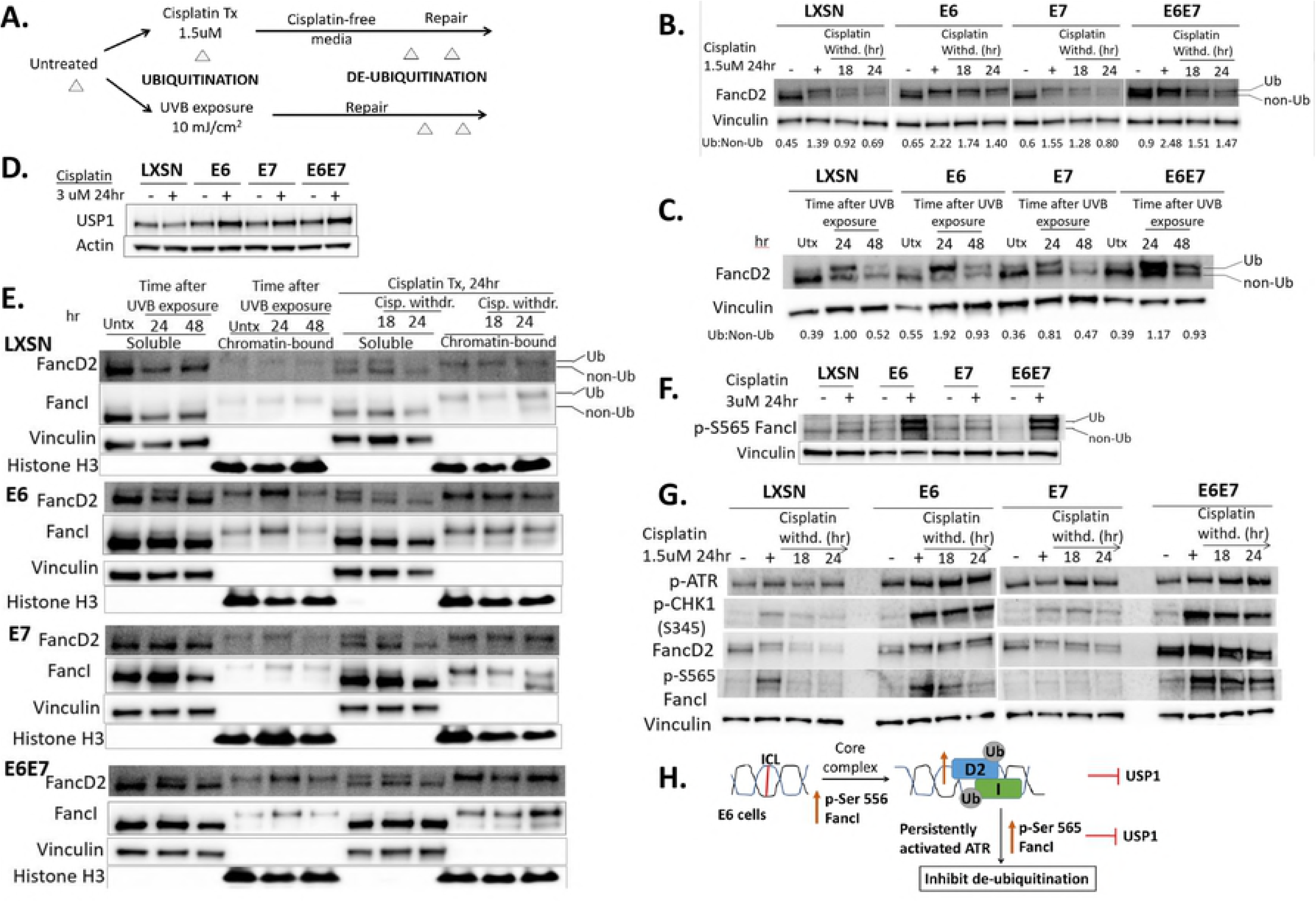
HPV E6 causes delayed de-ubiquitination of FancD2. (A) Outline of an experiment to evaluate FancD2 monoubiquitination/de-ubiquitination pattern. Transduced HFK cells were untreated or treated with cisplatin (1.5 uM for 24 hr) or exposed to 10 mJ/cm^2^ UVB and allowed to repair. Whole-cell lysates were prepared for immunoblot at various time points during the experiments (represented by Δ). (B-C) Immunoblots of HFKs subjected to the experiment outlined in Fig 4A, following cisplatin withdrawal (B) or recovery after UVB exposure (C). Ratios of monoubiquitinated to non-ubiquitinated FancD2 (D2 Ub: non-Ub) are indicated beneath the corresponding lanes. (D) USP1 immunoblot in cells untreated and treated with cisplatin. (E) Immunoblot of soluble and chromatin-bound fractions prepared from transduced HFK cells subjected to the experiment outlined in Fig 6A. Vinculin and Histone H3 act as loading controls respectively for soluble and chromatin-bound fractions. (F) Immunoblot showing levels of p-FancI-S565 in cisplatin-treated or untreated cells. (G) Immunoblot for p-ATR, pCHK1, FancD2 and p-FancI-S565 following cisplatin withdrawal for 18 and 24 hrs. Actin or vinculin act as a loading control for immunoblots (B-G). (H) Proposed mechanisms for delayed de-ubiquitination of FancD2 in E6 cells.

As Ub-FancD2 persists abnormally in E6 cells, a series of experiments were designed to determine the basis of the delay in de-ubiquitination. First, we asked whether this phenotype was a consequence of decreased levels of the FancD2 deubiquitinating enzyme, USP1. In fact, the opposite result was obtained: E6/E7 expressing cells showed elevated USP1 with cisplatin treatment, while basal levels of USP1 did not differ among untreated cells (Fig 5D). These results suggest that delayed FancD2 de-ubiquitination in E6 cells is not because of lower USP1 expression. Second, a recent study indicated that de-ubiquitination of FancD2/FancI by USP1 occurs only when the complex is no longer bound to chromatin. (45). Delayed FancD2 de-ubiquitination in E6 expressing cells may be due to retaining a chromatin-bound conformation of FancD2/FancI after DNA damage. To address this possibility, the association of FancD2/FancI with chromatin was examined using cell fractionation following cisplatin withdrawal or UVB exposure (Fig 5E). 24 or 48 hr following UVB exposure, E6 or E6+E6E7 expressing cells showed increased chromatin-bound Ub-FancD2/FancI, whereas LXSN and E7 expressing cells showed predominantly soluble non-Ub FancD2/I. There was also increased chromatin-bound Ub- FancD2/FancI at 18 and 24 hr following cisplatin treatment in E6 expressing cells compared to LXSN control. These data suggest that the delayed de-ubiquitination pattern observed in E6 expressing cells was due to FancD2/FancI being retained in a chromatin-bound conformation where USP1 cannot work efficiently to de-ubiquitinate the complex, despite high levels of USP1.

Third, while FancI phosphorylation at Serine 556 promotes FANCD2 monoubiquitination, its phosphorylation at Serine 565 inhibits FANCD2 de-ubiquitination and impairs ICL repair (7). Therefore, we investigated whether delayed de-ubiquitination of FancD2 was also due to elevated levels of phosphorylated-S565 FancI in E6 expressing cells. As expected, E6 or E6+E7 expressing cells showed increased p-S565 FancI on cisplatin treatment (Fig 5F). As phosphorylation of FancI occurs through the ATR-mediated pathway (7) and ATR/CHK1 activation increases FancD2 monoubiquitination (9), delayed FancD2 deubiquitination in E6 cells may be due to persistently activated ATR. Therefore, we examined the levels of p-ATR, p- S345 CHK1, p-S565 FancI as well as FancD2 monoubiquitination/deubiquitination patterns in cells which had recovered for 24 hr in normal media after cisplatin withdrawal. In E6 or E6+E7 expressing cells, ATR/CHK1 was activated and persisted following cisplatin withdrawal, compared to LXSN (Fig 5G). These results are consistent with persistent levels of both ubiquitinated FancD2 and S565-phosphorylated FancI in E6 expressing cells following cisplatin release. Further, persistence of pATR and pCHK1 foci following UV exposure was seen in E6 cells compared to LXSN (data not shown). These data demonstrate that persistent activated ATR/pCHK1/pFancI signaling could contribute to the delayed de-ubiquitination of FancD2 in E6 cells. Fig 5H shows the potential mechanisms for delayed de-ubiquitination of FancD2 in E6 cells.

### E6 does not abrogate the cell-cycle specific process of FancD2 monoubiquitination or de-ubiquitination

To determine whether the effect of E6 on increased monoubiquitination and delayed de-ubiquitination of FancD2 was a consequence of aberrant cell cycle progression, LXSN and E6 cells were synchronized in early S-phase by double-thymidine block, released, and FancD2 mono- or de-ubiquitination patterns were examined as cells progressed through the cell cycle (Fig 6). A previous study reported that FancD2 undergoes monoubiquitination during the S-phase of the cell cycle (4). Consistent with this study, in both LXSN and E6 cells, monoubiquitinated FancD2 was the major isoform present upon release from double-thymidine arrest into G1/S border and S-phase. Ub-FancD2 was noticeable till late S-phase, just prior to the decline of cyclin A. It is known that USP1 deubiquitinates FancD2 when cells exit S phase (12). Ub-FancD2 was de-ubiquitinated when both LXSN and E6 cells exited S phase. Cell-cycle analysis by flow cytometry was performed at each time point to track cell-cycle progression (Fig 6D). These data suggest that E6 does not abrogate cell cycle-specific process of FancD2 monoubiquitination and de-ubiquitination.

**Fig 6.**
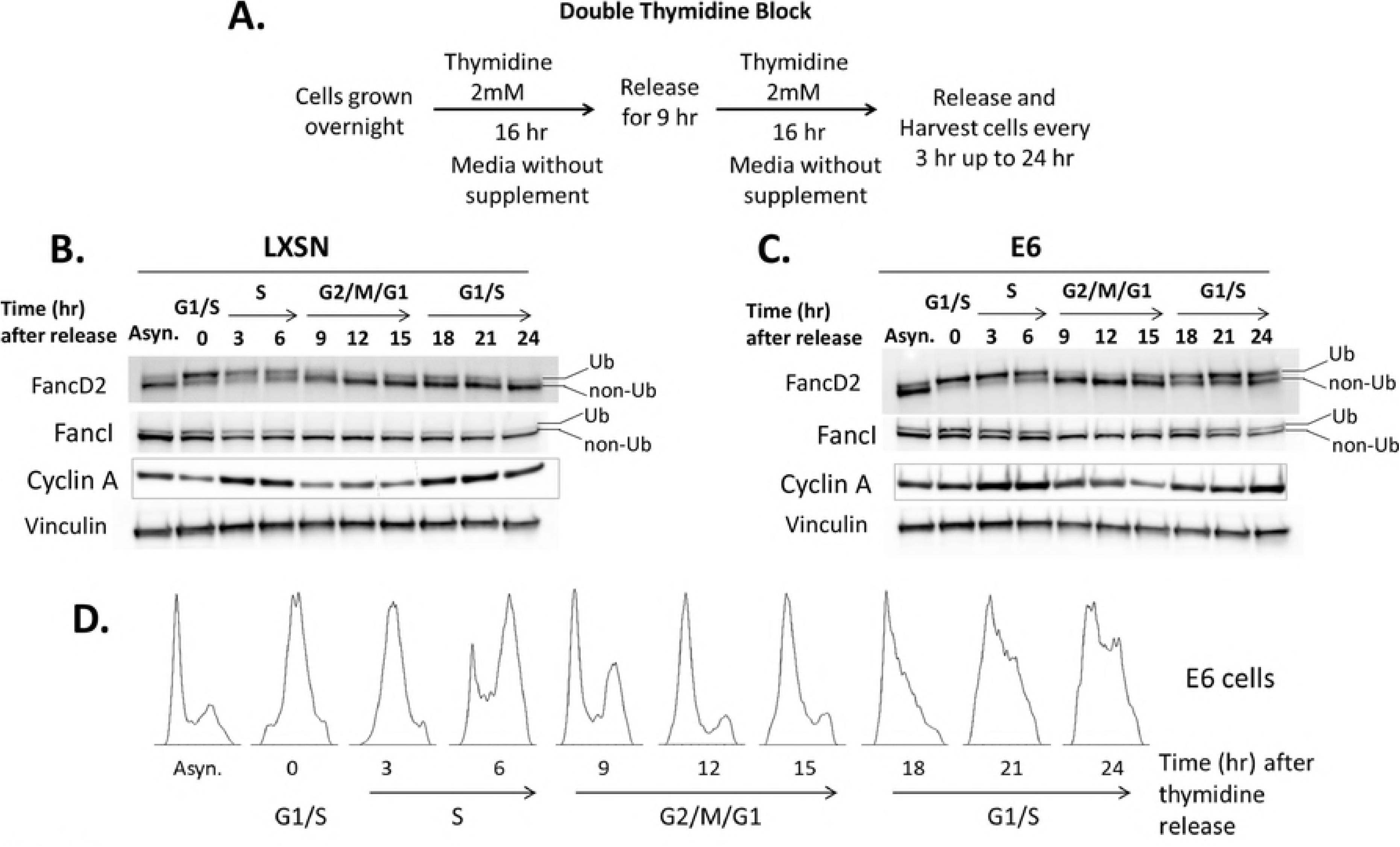
Cell-cycle specific process of FancD2 monoubiquitination/ de-ubiquitination is not abrogated in E6 cells. (A) Outline of synchronization assay in HFK cells using double-thymidine block. Cells were synchronized by double-thymine block and released at various time points. Immunoblot for FancD2, FancI, cyclin A and vinculin of LXSN (B) and E6 cells (C). Cell-cycle phases at each time point were determined by flow cytometry of DNA content (D). Asynchronous (Asyn.) cells, which have not undergone thymidine block, were included for comparison.

### E6 mediated increased FancD2 monoubiquitination is p53 independent

To further address the regulation of FancD2 mono or de-ubiquitination, the potential role of p53 was examined. The p53 tumor suppressor is a major cellular target of HPV16 E6. One study reported that p53 downregulates FancD2 mRNA and protein levels (46). As E6 degrades p53, we expected to see increased FancD2 mRNA and protein levels in E6 expressing cells. However, no significant differences in FancD2 mRNA levels between LXSN and E6 cells were observed (data not shown). But, since there was increased protein levels of total FancD2 in E6 expressing cells (Fig 3A), we hypothesized that this phenotype was dependent on the ability of E6 to degrade p53. To test this, HFK cells expressing a mutant of HPV16 E6 (8S/9A/10T) that is incapable of degrading p53 (47, 48) was used. Western blot analysis confirmed that E6 expression degrades p53, but the mutant failed to induce p53 degradation (Fig 7A). Both cell lines were also evaluated for E6 expression by RT-PCR (data not shown). Total FancD2 level was similar in mutant E6 and wild-type E6 cells (Fig 7B), indicating that E6 increased FancD2 protein level independently of its effects on p53 degradation. To confirm these results, cells were treated with Nutlin-3a, a Mdm2 inhibitor that increases p53 activity by preventing Mdm2-mediated proteasomal degradation. Nutlin treatment decreased total FancD2 level in LXSN cells but not in E6 expressing cells (Fig 7C). Similar downregulation of FancD2 was observed in lung fibroblast cells upon Nutlin-induced p53 activation, with no significant change in FancD2 level in p53-deficient counterparts (46).

**Fig 7.**
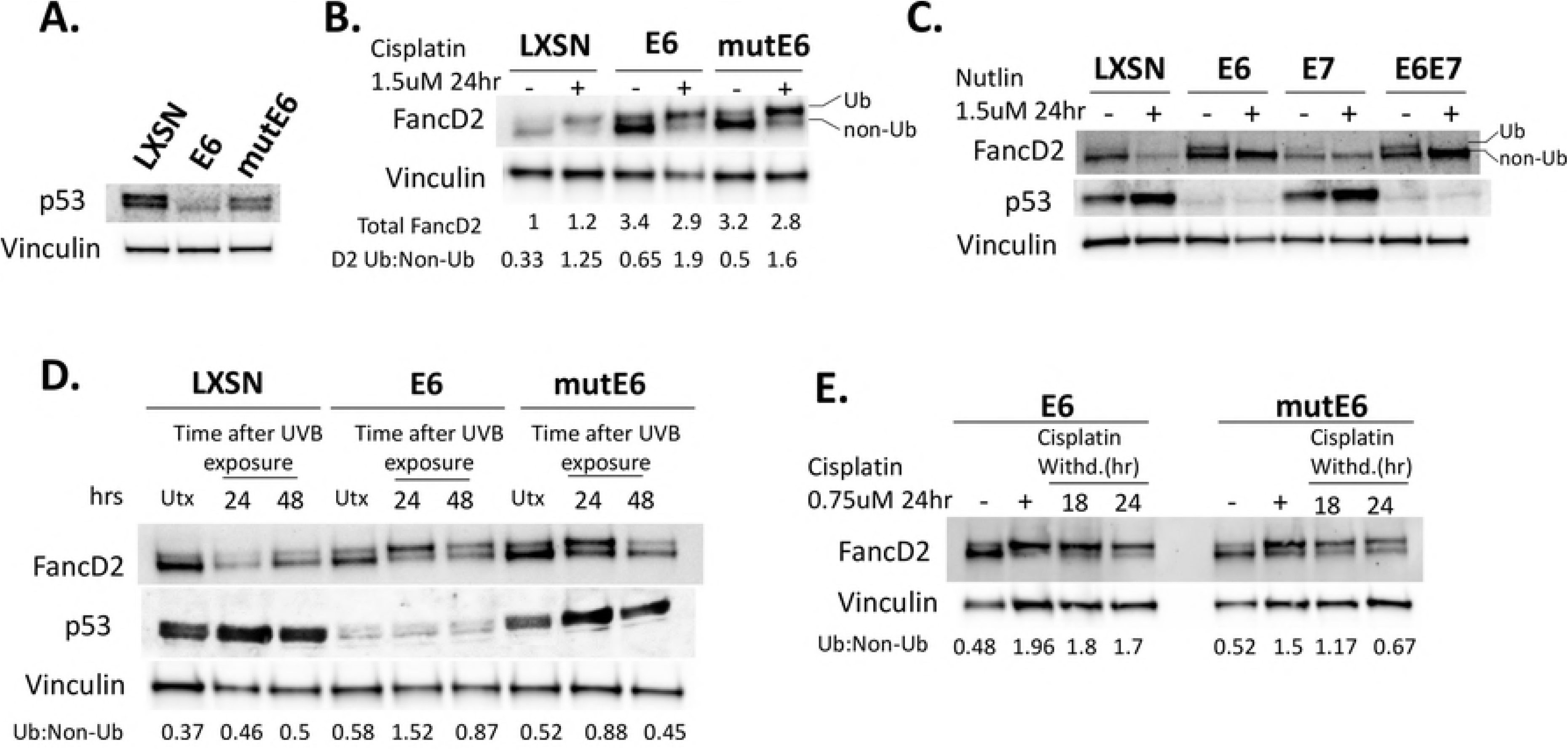
E6 mediated increased FancD2 monoubiquitination is p53 independent, but delayed FancD2 de-ubiquitination is dependent on p53 degradation. (A) Immunoblot for p53 in HFK cells transduced with LXSN, E6 and E6 mutant (8S/9A/10T). (B-C) Immunoblot showing FancD2 expression and monoubiquitination status in cells which were either untreated or treated with cisplatin (B) or nutlin (C) at 1.5 uM for 24 hr. (D-E) Cells were untreated or treated with cisplatin (E) or exposed to 10 mJ/cm^2^ UVB (D) and processed similarly as in Fig 5. See also Fig S6A.

Several publications suggested that FancD2 monoubiquitination is p53-independent. Several p53 defective cancer cell lines and chicken B lymphocyte cells lacking functional p53 were found to be fully competent for FancD2 monoubiquitination (49-51). Interestingly, Rego et al. showed that FancD2 mono-Ub is p53-independent but dependent on p21, its downstream target (49). In our study, although E6 showed increased Ub-FancD2 compared to LXSN, this increment was similar in mutant E6 cells, which fail to degrade p53 (Fig 7B). Hence, increased monoubiquitination of FancD2 by E6 is not a direct consequence of p53 degradation. To confirm these results, we used p53 knockdown LXSN cells and found that the Ub-FancD2 level was unchanged when compared to p53-sufficient LXSN (Fig S4).

### E6 mediated delayed FancD2 de-ubiquitination is dependent on p53 degradation

The exact role of p53 in FancD2 de-ubiquitination is not well-understood, although a study reported that p21, a p53 downstream target, represses USP1 transcription (49). Since E6 increased USP1 protein expression on cisplatin treatment (Fig 5D), we argued that the mechanism by which E6 causes delayed FancD2 de-ubiquitination may be related to its effects on p53 degradation. To examine this, we performed similar experiments as in Fig 4A in cells expressing the mutated E6 that is incapable of degrading p53. At 48hr after UV exposure, FancD2 de-ubiquitination pattern in mutant E6 was similar to LXSN control cells (Fig 7D, lanes 3 and 9). Similarly, after 24hr release from cisplatin treatment, mutant E6 showed de-ubiquitinated FancD2 (Ub: Non-Ub ratio of 0.67), whereas wild-type E6 cells had predominately monoubiquitinated FancD2 (ratio of 1.7) (Fig 7E). This indicates that delayed FancD2 de-ubiquitination in E6 cells was related to the ability to degrade p53. To confirm that the observed effects were specific consequences of p53 degradation, we used shRNA to stably knockdown p53 in LXSN cells and analyzed FancD2 Ub/de-Ub pattern upon UV exposure or cisplatin withdrawal. Once again, FancD2 de-ubiquitination was markedly delayed in the absence of p53 (Fig S5B). These results strongly support a p53-dependent effect of E6 in causing delayed FancD2 de-ubiquitination.

We next examined whether p53 dependent delayed FancD2 deubiquitination in E6 cells is due to activated ATR/CHK1/pFancI signaling. pATR/pCHK1 levels were reduced in cells expressing mutant E6 that cannot degrade p53 compared to wild-type E6 cells (Fig S5C). Reduction in ATR activity resulted in decreased phosphorylation of FancI at S565 and increased FancD2 de-ubiquitination in mutant E6 cells.

### E6 attenuates Palb2 expression and foci formation

To further understand how E6 or E7 impair the FA pathway and cause genomic instability, the expressions of proteins downstream of FancD2/I complex were evaluated. As a key regulator of the FA pathway, FancD2 recruits and coordinates functions of downstream FA repair proteins (52), including BRCA2/FancD1, BRCA1/FancS, Rad51/FancR, Palb2/FancN and BRIP1/FancJ. Previously, we demonstrated that neither BRCA2, BRCA1 nor Rad51 protein expression is impaired by HPV oncogenes (3). To determine whether the abundance of any other downstream protein is affected by HPV oncogenes, the levels of Palb2 were measured. Palb2 level was reduced when E6 was expressed separately or together with E7 (Fig 8A). Depleted Palb2 level in E6 cells was not a consequence of p53 degradation (Fig S6). Because E6 reduced Palb2 levels, the ability of Palb2 to form nuclear foci in response to cisplatin was examined. In LXSN control cells, the percentage of cells with >5 Palb2 foci peaked at about 90% following cisplatin treatment. In contrast, significantly fewer (~60%) E6 expressing HFKs formed Palb2 foci on cisplatin treatment (Fig 8B). Although fewer Palb2 foci were observed in E6 cells, these foci perfectly localize to pH2AX in I-SceI transfected U2OS-DR system (Fig S7A). Because E6 reduces Palb2 expression and foci formation, and Palb2 is also a component of FA nuclear foci for ICL repair, we asked whether the cisplatin hypersensitivity observed in E6 expressing cells is due to a defect in Palb2. Genetic epistasis analysis for cisplatin sensitivity was conducted by knocking down Palb2. Palb2 depletion in LXSN and E7 cells resulted in further sensitization to cisplatin (Fig 8C), whereas there was no effect on cisplatin sensitivity in E6 and E6+E7 expressing cells. These results suggest that cisplatin sensitivity seen in E6 expressing cells is, in part, due to a defect in Palb2 expression and foci formation.

**Fig 8.**
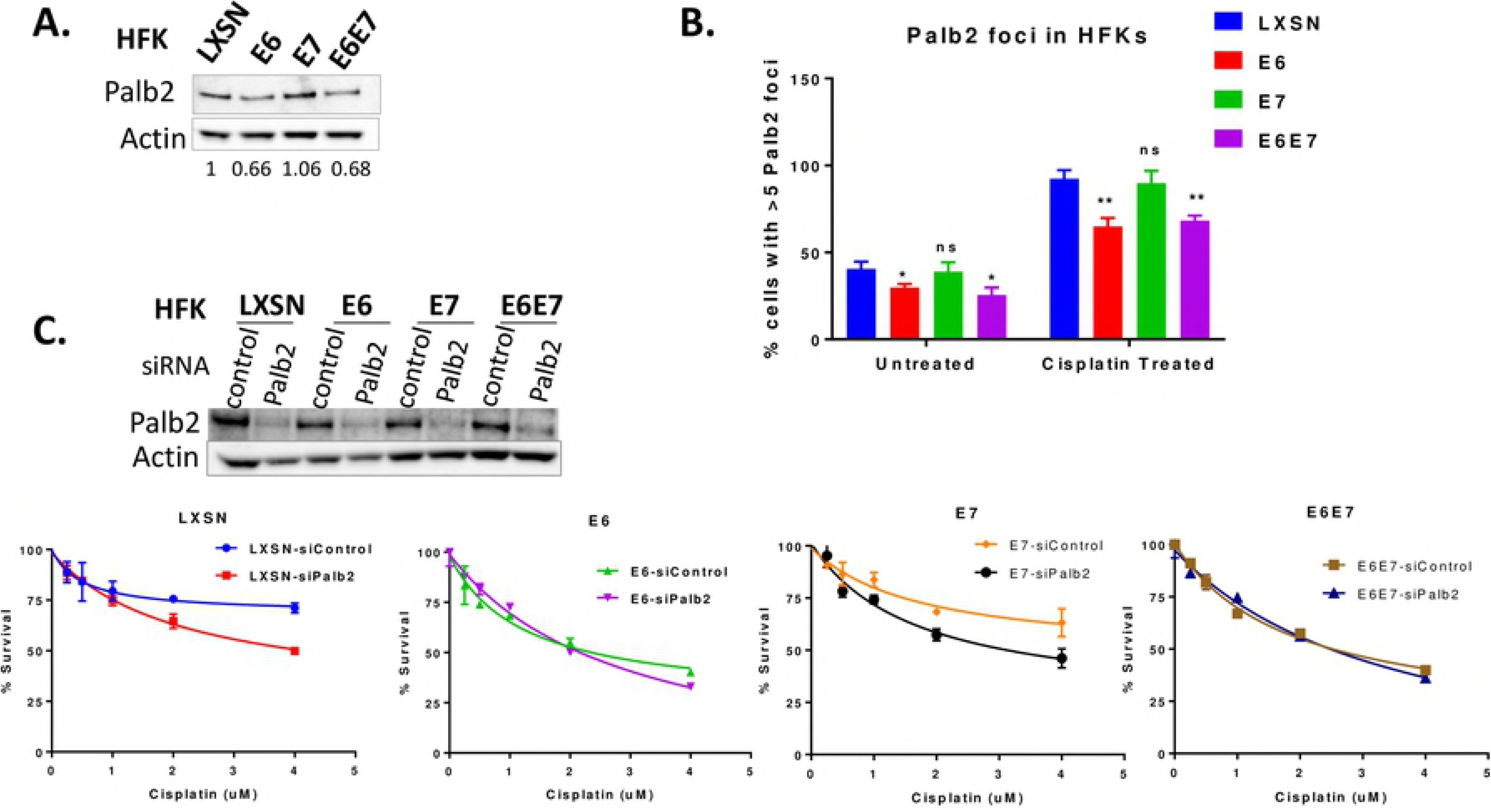
E6 attenuates Palb2 expression and foci formation. (A-C) HFKs were transduced with LXSN, E6 or E6/E7 (A) Immunoblot showing Palb2 expression. Actin serves as a loading control. (B) Cells were untreated or treated with cisplatin (3 uM) for 24 hr and immunostained with Palb2, pH2AX, and DAPI. Cells with >5 Palb2 foci were counted and the percentage of positive cells is plotted (n = 3, mean ± SD). * and ** indicate significance respectively at p<0.05 and p<0.01 (compared to LXSN) whereas ns indicates non-significant. (C) HFKs were transfected with siControl or siPalb2. Immunoblot confirming depletion of Palb2 (upper panel). Cisplatin survival assay for siRNA-transfected cells (lower panel).

Reduced Palb2 expression and foci formation in E6 cells (Fig 8A-B) may affect the interaction of Palb2 with other FA repair proteins, such as Rad51, required for resolution of ICLs. In fact, Palb2 physically and functionally interacts with Rad51. First, Palb2 colocalizes with BRCA2 foci, which also includes Rad51 (53-56). Second, loss of Palb2 impairs Rad51 focus formation (57) and there is Palb2-dependent loading of BRCA2-Rad51 repair machinery at DSBs (56). Therefore, we next examined whether the defect in localization of Rad51 observed in E6 expressing cells is due to decreased Palb2 expression or foci formation. In all HFK cells, including E6 expressing cells, Palb2 depletion caused a dramatic reduction in colocalization of Rad51 to pH2AX in I-SceI induced DSB (Fig S7B) indicating that the Rad51 recruitment defect observed in E6 cells is not due to impaired Palb2 expression and foci formation.

## DISCUSSION

The FA DNA repair pathway is involved in guarding genome integrity, especially when challenged by endogenous and exogenous DNA crosslinking agents. Cisplatin is the most commonly used chemotherapeutic drug for cervical and oropharyngeal cancers. Here we provide mechanisms by which HPV oncogenes attenuate the FA pathway and contribute to tumorigenicity and the response to cisplatin therapies in HPV-associated malignancies. Our present study (Fig 1) along with others (24) confirm the cross-linker hypersensitivity in HPV oncogene expressing cells. We also show that E6/E7 expressing cells have a defect in repairing cisplatin-induced ICLs (Fig 2). The epistatic analysis confirmed that ICL sensitivity in E6/E7 cells was due to a defect in FancD2 or downstream of FancD2 (Fig 1). However, FancD2 monoubiquitination and foci formation were increased in E6 or E7 cells (Fig 3-4), which is consistent with the previous work conducted in HPV+ and HPV oncogene expressing cells (27, 31). Strikingly, Ub-FancD2 was seen without cisplatin treatment in E6 expressing cells, which might be due to the presence of activated ATR which increases FancD2 monoubiquitination (9). HPV oncogenes though, promoted the initiation of FA pathway by increasing FancD2 monoubiquitination and foci formation but hindered the completion of the repair by multiple mechanisms. E6 or E7 caused the accumulation of monoubiquitinated FancD2 at sites away from DNA damage. This subsequently hindered the recruitment of downstream protein Rad51 to DNA damage sites in E6 cells (Fig 4D-H). Further, E6 mediated p53 degradation does not hamper cell cycle specific process of FancD2 modifications but abrogate the repair by delaying FancD2 de-ubiquitination (Fig 5-7). In addition, E6 reduced the expression and foci formation of Palb2 (Fig 8). Though there was a defect in FancD2 localization to DSBs in E6 cells, no mislocalization of Palb2 was observed, which is in support of a recent study reporting that FancD2 and Palb2 localize to DSBs independently of each other (58).

HPVs have been shown to recruit numerous cellular repair factors to their replication centers, mainly for HPV amplification (59-61). We speculate that increased Ub-FancD2 and foci formation may create an environment where FA repair proteins are readily available to support HPV replication. Consistent with our speculation, a study showed that higher levels of FancD2 and larger foci are predominantly recruited to HPV DNA rather than cellular genomes and localize to viral replication centers (31). The preferential recruitment of cellular repair proteins to HPV replication centers would facilitate viral replication but could leave chromosomal DNA unrepaired. Our present work shows how HPV delays the repair of ICLs and attenuates the FA DNA repair pathway. Mislocalization of FancD2 observed in our study may represent a fruitless attempt to bring the FancD2 repair protein to non-existing viral replication centers. This would attenuate ICL repair regardless of the presence or absence of the viral genome because FancD2 helps in the recruitment of downstream FA repair proteins, including Rad51 in nuclear foci at the site of DNA damage. This is supported by our data demonstrating that mislocalized FancD2 in E6 cells causes a reduction in Rad51 recruitment to DSBs (Fig 4G-H).

FancD2 monoubiquitination is necessary for ICL repair, but, by itself, is not sufficient for an efficient FA pathway. In other words, having more monoubiquitinated FancD2 within a cell does not necessarily improve ICL repair. This is exemplified by studies demonstrating that USP1 -mediated FancD2 de-ubiquitination is required for both FancD2 foci formation and ICL sensitivity (10, 13) and for a functional FA pathway (7, 12, 45). HPV E6 caused the delayed de-ubiquitination of FancD2, impairing ICL repair (Fig 5). HPV E6 cells showed increased phosphorylated S565 FancI and persistently activated ATR/CHK-1, which may, in part, cause delayed FancD2 de-ubiquitination in these cells. The delayed de-ubiquitination pattern observed in E6 expressing cells may be also due to a block at the step of release of FancD2 from chromatin which hinders USP1 deubiquitination activity.

ATR/CHK1 signaling is a key step in activating the FA pathway (5). ATR is activated when the replication fork encounters DNA damage. ATR then phosphorylates several substrates, including FancD2 and FancI as well as members of the FA core complex. ATR-mediated FancI phosphorylation at Serine 556 (called ubiquitination-independent phosphorylation) occurs predominantly upstream of, and promotes, the monoubiquitination of FancD2/FancI (7, 62). On the other hand, FancI Serine 565 phosphorylation (called ubiquitination-linked phosphorylation) occurs downstream of Ub-FancD2/I and inhibits FancD2 de-ubiquitination (7). Augmented ATR activity in E6 expressing cells resulted in increased phosphorylation of FancI at both sites S556 and S565. Increased p-S556 FancI promotes the monoubiquitination of FancD2, but at the same time elevated p-S565 FancI inhibits de-ubiquitination.

Together, we provide a mechanistic framework for HR-HPV oncogenes disruption of the FA DNA repair pathway (Fig S8). This study advances our understanding of HPV tumorigenesis as well as tumor sensitivity to cisplatin during chemotherapy. Our study suggests a general model for the progression of HPV associated cancers. Over the multi-decade course of HPV-induced tumorigenesis, persistent expression of HPV oncogenes impairs the FA pathway. This leads to cells with destabilized genomes, allowing for rapid tumor progression. Previous studies are in support of the model that dysregulated FA pathway contributes to the development of cancers in patients without FA (63-65) and with FA (66, 67). An extremely high incidence of cancer in FA patients (68) further suggest that the inactivation of FA pathway results in tumor progression.

Our work also has important therapeutic implications. Because continued expression of HPV oncogenes is required for cancer development, most HPV-associated tumors likely have defective FA pathway. These defects result in cisplatin hypersensitivity and demonstrate the mechanisms underlying the therapeutic efficacy of cisplatin in HPV-associated cancers. This also explains the reason underlying the better response rates of HPV+ oropharyngeal cancers than HPV negative head and neck cancers to cisplatin treatment. We believe that tumors that are cisplatin resistant, may have adapted other unexplored mechanisms to escape these defects. One possibility is that some cells with an intact FA pathway are positively selected during cancer progression, resulting in the growth of a cisplatin-resistant tumor.

## EXPERIMENTAL PROCEDUTES

### Ethics Statement

The use of deidentified neonatal human foreskins for the study was approved by the Institutional Review Board at the Swedish Medical Center (Seattle, WA).

### Cell Culture, transduction, and treatment

Primary human foreskin keratinocytes (HFKs) were generated from deidentified neonatal human foreskins and grown in EpiLife Medium supplemented with 60 μM calcium chloride (ThermoFisher Scientific, MEPI500CA) and human keratinocyte growth supplement (ThermoFisher Scientific, S0015). HFKs derived from multiple donors were used to repeat the results. U2OS DR-GFP cells (a gift from Maria Jasin) were grown in DMEM supplemented with 10% FBS. U2OS DR-GFP cells contain a single integrated copy of the DR-GFP cassette (69).

Retrovirues were produced in 293T cells (Thermo Fisher Scientific) using plasmid constructs (LTR/VSV-G, CMV/tat, pJK3 and pLXSN vector-empty, E6, E7 or E6E7, or E6 8S/9A/10T mutant) and TransIT-293 transfection reagent (Mirus Bio) according to manufacturer’s protocol. Following retroviral transduction and G418 selection (50ug/ml) in HFKs or U2OS-DR cells, the expression of HPV16 oncogenes E6 and E7 was confirmed by qRT-PCR of 16 E6 and E7 and immunoblot to p53 and pRB (well-characterized cellular targets). The expression of p53 in HFKs was suppressed using transduction-ready p53 shRNA lentiviral particles (Santa Cruz Biotechnology, sc-29435-V). HFKs were transduced with lentiviral stock in media containing 10ug/ml polybrene. Following 48 hr of transduction, stably transduced cells were selected in 0.5 ug/ml puromycin and the efficiency of knockdown was monitored using immunoblot to p53.

Cells were treated with cisplatin (Selleck Chemicals, S1166), Mitomycin C (MMC) (Sigma, M4287), Nutlin-3a (Selleck Chemicals, S8059), Ionizing radiation (IR) (GammaCell Cesium Irradiator −1000) or UVB (two FS20t12/UVB bulbs, Solarc Systems, Inc.). Cisplatin was dissolved at the stock of 1.5mM in 0.9% NaCl (sterile), Nutlin-3a at the stock of 10mM in DMSO, aliquoted and kept at −20° C protected from light.

### Antibodies

Primary antibodies against p53 (Cell Signaling Technology, 9282), pRb (BD Pharmingen, 554136), FancD2 (Santa Cruz Biotechnology, FI17, sc-20022 or Abcam, ab2187), FancI (Santa Cruz Biotechnology, A-7, sc-271316), Phospho-Ser556 FancI and Phospho-Ser565-FancI (gifts from Ronald Cheung, Toshiyasu Taniguchi lab), pH2AX (Millipore, 05-636), Rad51 (Cosmo Bio Co. Ltd, 70-001), Palb2 (Bethyl Laboratories, A301-246A), Palb2 (a gift from Bing Xia, Rutgers Cancer Institute), Phospho-ATR (Ser428) (Cell Signaling Technology, 2853), Phospho-Chk1 (Ser345) (133D3) (Cell Signaling Technology, 2348), PCNA (PC10) (Cell Signaling Technology, 2586), Ubiquityl-PCNA (Lys164) (Cell Signaling Technology, 13439), UHRF1 (Santa Cruz Biotechnology, H-8, sc-373750), USP1 (C-term; a gift from Tony Huang, NYU School of Medicine), Cyclin A (in-house, Clurman lab), Beta actin (GeneTex, GTX110564 or Cell Signaling Technology, 5125), Vinculin (Sigma, V9131), and Histone H3 (Abcam, ab1791) were used. Secondary antibodies were conjugated with horse radish peroxidase (mouse or rabbit, Cell Signaling Technology) or appropriate Alexa Fluor (Molecular Probes) were used.

### Cell extraction, Fractionation, and Immunoblotting

Whole cell lysates were prepared by directly lysing cell pellets in SDS sample buffer (0.05 M Tris-HCl, (pH 6.8), 2% SDS, 6% β-mercaptoethanol) and boiling for 5 min. Cell fractionation was performed to obtain soluble and chromatin-bound fractions, as described (70). Briefly, cell pellets were re-suspended in CSK + 0.1% Triton-X buffer (10mM PIPES, pH = 6.8, 100mM NaCl, 1mM EGTA, 1mM EDTA, 300mM Sucrose, 1.5mM MgCl2, 0.1% Triton-X-100 and protease inhibitors) and incubated on ice for 5 mins. After centrifugation (1500g for 5 min), the supernatant was collected and stored (soluble fraction). Pellet (Chromatin-bound fraction) was washed twice in CSK buffer and then resuspended in SDS sample buffer and boiled for 5 min. Samples were stored at −20 °C until quantification and immunoblotting. Proteins were quantified using Bradford protein assay (Bio-rad) and processed using NuPAGE LDS sample buffer (ThermoFisher Scientific) for loading equal amounts (15ug) onto the gels. SDS-PAGE electrophoresis was done using NuPAGE 3–8% Tris-acetate or 4–12% Tris-glycine gels (ThermoFisher Scientific) and proteins were transferred to Immobilon-P PVDF membranes (Millipore). Primary antibodies were diluted in blocking buffer (4% milk in TBS-Tween 20) or in 3% BSA in TBST for phosphorylated proteins and incubated overnight at 4 °C. Horseradish peroxidase (HRP)-conjugated anti-mouse or anti-rabbit secondary antibodies were used. Blots were developed using Clarity Western ECL substrate (Bio-Rad) and images were acquired using ChemiDoc MP Imaging System (Bio-Rad) and processed using Image Lab Software (Bio-Rad). Band quantification was performed using ImageJ, normalizing with a loading control.

### RT-PCR

qRT-PCR was conducted to evaluate the expression of HPV16 E6, E7, and GADPH genes, as described previously (3). To evaluate the expression of FancD2 mRNA, two sets of primers were designed to amplify the longer FancD2 isoform from transcript sequence NM_033084.4. The first primer set adapted from Jaber et al. (46) was Forward 5’-AGACTGTCAAAATCTGAGGATAAAGAGA-3’ and Reverse 5’-TGGTTGCTTCCTGGTTTTGG-3’; and next set was designed as Forward 5’- CATGGCTGTTCGAGACTTC-3’ and Reverse 5’-CACAAAGAGACGCCCATAC-3’.

### Cisplatin Cell Survival Assay

Cell survival over a range of cisplatin concentrations was measured using a crystal violet absorbance-based assay (71). Briefly, cells were seeded on 24-well plate overnight and treated with cisplatin at indicated doses for 3 days. Cells were then washed once with PBS and fixed for 10 min at room temperature in 10% methanol and 10% acetic acid. Adherent cells were stained with 0.5% crystal violet in methanol. Plates were rinsed in water and allowed to dry completely. The adsorbed dye was re-solubilized with methanol containing 0.1% (wt/vol) SDS by gentle agitation for 2 hr at room temperature. Dye solution (200 μl) was transferred to 96-well plates and diluted (1:2) in methanol. Absorbance was measured at 595 nm. Cell survival was calculated by normalizing the absorbance relative to untreated controls.

### siRNA transfection

siRNAs were transfected at a final concentration of 20nM in 6-well or 10 cm plates using Lipofectamine RNAiMAX (Invitrogen), following the manufacturer’s Reverse Transfection protocol. I-SceI colocalization and crystal violet assays were conducted, respectively 24 and 48 hr after siRNA transfection. The following siRNAs were used: si-FancD2: 5’- AACAGCCAUGGAUACACUUGA-3’; si-Palb2 (Santa Cruz Biotechnology, sc-93396) and siRNA Universal Negative Control#1 (Sigma, SIC001). Knockdown was confirmed by immunoblotting.

### Immunofluorescence Microscopy

HFKs and U2OS DR-GFP cells (transduced with LXSN, 16 E6, 16 E7 and 16 E6E7) were grown in chamber slides overnight. Cells were either left untreated or treated with cisplatin or transfected with I-SceI expression vector (Addgene, 26477). Cells were then washed with 1X PBS and fixed and co-permeabilized using 2% paraformaldehyde and 0.5% Triton-X in PBS for 30 mins. After washing, cells were blocked with 3% BSA and 0.1% Tween20 in PBS for 1 hr. Cells were then subsequently incubated with appropriate primary antibodies (indicated in the text) in a moist chamber at 4° C overnight. The following day, cells were washed 3 times with 1X PBST (1X PBS + 0.1% Tween20) for 5 min each and then incubated for 1 hr at room temperature with Alexa Fluor secondary antibodies in dark. Primary and secondary antibodies were diluted in blocking buffer. Following secondary antibody incubation, cells were washed and mounted using coverslip in ProLong™ Diamond Antifade Mountant with DAPI (Molecular Probes). Cells were then visualized using a DeltaVision Elite confocal microscope (Applied Precision). The images were deconvolved using the Deltavision SoftWoRx program and were analyzed using ImageJ. Each experiment was repeated at least 3 times independently.

### I-SceI Colocalization assay

I-SceI colocalization assay in U2OS-DR cells (transduced with LXSN, E6, E7 or E6E7) were performed as described previously (3). Briefly, cells were transfected with I-SceI expression vector using TransIT-LT1 reagent (Mirus) and were stained for pH2AX and the indicated protein. In the case of gene knockdown experiments, reverse siRNA transfection was done a day before transfecting I-SceI plasmid. A single large pH2AX foci created by I-SceI expression was inspected for its colocalization with indicated repair protein.

### Alkaline Comet Assay

Repair of interstrand crosslinks was assessed by an alkaline comet assay (32-34) using Trevigen’s CometAssay (4250-050-K), following the manufacturer’s protocol with some modifications. Cells were treated with 1.5 uM cisplatin for 2 hr. At the end of treatment, cells were washed with PBS and incubated in fresh medium for the required post-incubation time or harvested immediately (time 0 h). To process all the samples concurrently and to eliminate assay variability, cells were harvested and cryopreserved in media containing 10% DMSO and 50% FBS, prior to performing comet assay. Immediately before analysis, cells were thawed in ice-cold DPBS and irradiated with 12 Gy gamma irradiation (using GammaCell Irradiator GC-1000) to introduce a fixed number of random DNA strand breaks and processed according to Trevigen’s alkaline Comet assay procedures. Briefly, irradiated or unirradiated cells were immediately plated in low melting agarose on slides. After hardening of the agarose gel, cells were lysed and subjected to alkaline electrophoresis for 30 min at 4° C. After air-drying, DNA was visualized using SYBR Gold and images were captured using TissueFAXS machine using FITC filter. Images were analyzed using OpenComet Plugin tool in NIH ImageJ (72). Default background correction was used. The output images resulting from the automatic analysis were manually reviewed to eliminate outliers and select uniform comets. Updated tail moment values (Mean ± SE from >350 comets) were then used to calculate % ICL remaining.

The degree of DNA ICLs present in the cisplatin-treated sample was calculated by comparing the tail moment of cisplatin-treated and irradiated (Cp.IR) samples with irradiated untreated (IR) samples and unirradiated untreated (Ø) control samples. The level of ICL is proportional to the decrease in the tail moment in the irradiated cisplatin treated sample compared to the irradiated untreated control. To quantify ICL repair (or the percentage of ICL remaining), we employed the following formula: % Decrease in tail moment (or ICL remaining) = [1- (Cp.IR – Ø) / (IR – Ø)] x 100 where Cp.IR = mean tail moment of cisplatin-treated and irradiated cells; Ø = mean tail moment of untreated and unirradiated cells, and IR = mean tail moment of irradiated and untreated cells. The data were expressed as the percentage of ICLs that remained at a specific time point where 0 hr was normalized to 100%.

### Cell-cycle synchronization and analysis

Cells were synchronized at G1/S phase boundary by double-thymidine block and cell pellets were harvested for immunoblotting and flow cytometry analysis as described previously (4, 73), with some modifications. Briefly, HFK cells were treated with 2 mM thymidine (Sigma-Aldrich) in EpiLife media without growth supplements for 16 hours. Cells were then washed twice with PBS and released/grown in thymidine-free complete EpiLife media for 9 hours. Thereafter, cells were treated again with 2 mM thymidine in growth supplement-free EpiLife media for another 16 hours. Cells were washed twice with PBS and then released in complete media and harvested every 3 hours after release. Synchronized cells were analyzed at different time points by immunoblotting and analyzing DNA content by DAPI staining. Approximately 10,000 cells were analyzed using flow cytometry (BD FACS-Canto II), and flow histograms were generated using FlowJo software.

### Statistics

All statistical analyses were done using Student’s t-test (Unpaired two-tailed) in GraphPad Prism. P value < 0.05 was considered significant. Statistical significance at P < 0.05, P < 0.01 and P < 0.001 are indicated as *, **, ***, respectively.

## ACKNOWLEDGMENTS

We acknowledge Ronald Cheung from the Taniguchi lab for helpful guidance, advice, and discussions. Ahmed Diab from Clurman lab was helpful in designing experiments and having helpful discussions. We thank Bing Xia (Rutgers Cancer Institute) for Palb2 antibody and Tony T. Huang (NYU School of Medicine) for the USP1 antibody. We acknowledge current and former members of Galloway lab for assistance and suggestions. We are grateful to Ann Roman for critical reading of this manuscript. We thank Julio Vazquez Lopez and David McDonald of the FHCRC Scientific Imaging Facility for their help in immunofluorescence imaging. Staff of the FHCRC Flow Cytometry Facility were very helpful with FACS analysis.

## Supporting Figures

**Fig S1. The Fanconi anemia repair pathway.** Endogenous (aldehydes) and exogenous (cisplatin) agents cause interstrand crosslink lesions (ICL). The stalled replication fork at ICL site activates ATR signaling and recruits the FA core complex (FancA, B, C, E, F, G, L, and M). FancL with the help of FancT catalyze the monoubiqiuitination of the FancD2-FancI (ID) complex. Monoubiquinated ID complex helps in the recruitment of downstream repair proteins, including FancD1/BRCA2, FancS/BRCA1, FancN/Palb2, FancR/Rad51, and FancJ/BRIP1. These proteins crosstalk with other DNA repair pathways such as the nucleotide excision, homologous recombination and translesion synthesis to repair ICLs. ATR-mediated FancI phosphorylation at Serine 556 occurs upstream of, and promotes, the monoubiquitination of ID complex, whereas phosphorylation at Serine 565 occurs downstream of monoubiquitination and inhibit FancD2 deubiquitination.

**Fig S2. E6 expressing cells showed high Ub-FANCD2 & -FANCI both at baseline and after cisplatin/ MMC treatment.** (A-B) Confirmation of HPV16 E6 and E7 expression by qRT-PCR (A) and immunoblot of p53 and pRb in HFK cells (B). Immunoblot showing FancD2/FancI expression and monoubiquitination status in LXSN and E6 cells which (C) were either untreated or treated with 60ng/ml mitomycin C for 24 hr, and (D) were exposed to 10 mJ/cm2 UVB and incubated for indicated time points. (E) Immunoblot of transduced HFK cells harvested following different lengths of cisplatin treatment. Ub refers to the monoubiquitinated forms of FancD2 and FancI, and non-Ub refers to the non-ubiquitinated forms. Asterisks (*) indicate a non-specific band.

**Fig S3. Rad51 colocalization with FancD2 in HFK-LXSN cells.** HFK cells (transduced with LXSN) were treated with cisplatin (3 uM for 24 hr) and immunostained with FancD2 (red), Rad51 (green) and DAPI (blue). Representative images are shown.

**Fig S4. P53 knockdown does not change total and monoubiquitinated levels of FancD2.** (A) Immunoblot showing p53 knockdown in or p53 shRNA cells compared to LXSN control. (B) Immunoblot showing FancD2 expression and monoubiquitination status in HFK LXSN and p53 knockdown cells which were either untreated or treated with 1.5 uM cisplatin for 24 hr.

**Fig S5. Delayed FancD2 deubiquitination in E6 cells is dependent on p53 degradation.** (A) Immunoblots showing FancD2 mono- and deubiquitination status in HFKs which were treated with 1.5 uM cisplatin (upper panel) or 0.75uM cisplatin (lower panel) for 24 hr and allowed to repair, following cisplatin withdrawal. Initial experiments treating mutE6 cells with 1.5uM cisplatin and subsequent withdrawal did not give a clear idea (the deubiquitination pattern was more likely in between LXSN and E6). Therefore, less concentrated cisplatin (0.75uM) was used (Fig 7E). However, in case of LXSN cells, 0.75uM cisplatin treatment for 24hr was not enough to induce predominant mono-ubiquitinated FancD2 fraction. (B) Immunoblots showing FancD2 mono- and deubiquitination status in LXSN and p53 knockdown cells following UVB exposure (upper panel) and cisplatin withdrawal (lower panel). (C) Immunoblot for p-ATR, pCHK1, FancD2 and p-FancI-S565 in E6 and mutant E6 cells following cisplatin withdrawal for 18 and 24 hrs. Vinculin acts as a loading control.

**Fig S6. E6-mediated decreased Palb2 level is p53 independent.** Immunoblot showing Palb2 expression in (A) HFKs transduced with LXSN or E6/E7 untreated or 2hrs after UV exposure (B) in HFKs transduced with LXSN, E6 or mutant E6. Vinculin serves as a loading control.

**Fig S7. E6 does not affect the colocalization of Palb2 to DSBs and Rad51 recruitment defect associated with E6 is not due to Palb2.** (A) U20S-DR cells transduced with LXSN, or E6/E7 were transfected with I-SceI expression plasmid to induce DSB. Representative images showing cells with a single large pH2AX focus (red), were inspected for the colocalization with Palb2 (green), upper panel. Quantification of the frequency of colocalization of Palb2 with pH2AX focus (right panel). Data represent mean ± SEM and was based on observations from ≥ 25 cells from 3 independent experiments. (B) U2OS-DR cells were transfected first with the siControl or siPalb2 and then with I-SceI plasmid and were stained with Rad51 and pH2AX antibodies. Representative images showing U20S E7 cells stained with pH2AX (red), Rad51 (green), and DAPI, upper panel. Graph showing %Rad51 colocalization in siPalb2 cells compared to similarly transfected cells (right panel). * and ** indicate significance respectively at p<0.05 and p<0.01 (compared to siControl cells)

**Fig S8. HPV oncogenes impair the Fanconi anemia repair pathway.** E6 or E7 causes persistence of DNA interstrand crosslinks and increases cisplatin sensitivity. HPV oncogenes though promote the initiation of the FA pathway by increasing FancD2 monoubiquitination and foci formation, they attenuate the completion of the pathway by multiple mechanisms. First, E6/E7 reduces the recruitment of FancD2 to double strand breaks (DSBs). Second, E6 causes FancD2 mediated mislocalization of Rad51 from DSBs. Third, E6 reduces Palb2 level and foci formation. Lastly, E6 causes delayed FancD2 deubiquitination, impairing the functional FA pathway.

